# Phenotypic analysis of the edible fruits of *Lardizabala biternata*, an endemic and monotypic vine from the Chilean biodiversity hotspot in South America

**DOI:** 10.1101/2024.06.11.598437

**Authors:** Jaime Herrera, Leonardo D. Fernández

## Abstract

**Introduction:** *Lardizabala biternata* is a vine endemic to Chile, distributed between 32°S and 40°S. Its sweet edible fruits have historically been harvested by hand from the wild as there are no productive systems for this vine. Herein, we conducted the first phenotypic analysis of *L. biternata* fruits, which includes qualitative and quantitative analyses of morphological and morphometric traits. This phenotypic analysis is the baseline for the development of production systems that could reduce anthropogenic pressure on wild populations and favour the *ex-situ* conservation of this vine.

**Materials and methods:** We collected 282 fruits from two geographically distant populations during four fruiting seasons. In all of them we recorded 14 morphological attributes, including total weight, length, width, height, diameter, volume, edible pulp content, seed number weight and individual seed weight. We investigated morphometric differences between populations and seasons by analysis of variance (ANOVAs), phenotypic correlations by regressions and associations between traits by principal component analysis (PCA).

**Results:** On overage, fruits weighed 20.8 g (3.0 – 44.6 g) and measured 54 mm in length (20.1 – 83.4 mm) and 23.7 mm in diameter. Edible pulp contributed around 44.4% of total fruit weight. Observed traits displayed significant variations between seasons and among traits (length vs width vs height). Fruit weight showed a high correlation with edible pulp weight, fruit length, seed weight, seed number, and others.

**Discussion and conclusions:** Our study represents the first phenotypic analysis of the fruits of this wild, endemic, and rare plant. We comprehensively describe the morphological and morphometric characteristics of its fruits. The characteristics of *L. biternata* fruits show significant morphometric variation between populations and seasons. However, the edible pulp consistently remains the main component of the fresh fruit weight. Like other domesticated members of the Lardizabalaceae, the fruits of this wild plant have the potential for cultivation through the development of sustainable production systems. The information we provide serves as a baseline for the development of such systems through selection and genetic improvement of the plant.

## Introduction

*Lardizabala* [1] is a monotypic genus of flowering plants in the family Lardizabalaceae. The only representative of this genus is the vine *Lardizabala biternata* Ruiz & Pav., an endemic species of Chile, commonly known as cogüilera, voqui-cóguil, coile, cógüil, among other names [1, 2]. In Chile, this vine is distributed in the Chilean Winter Rainfall-Valdivian Forest global hotspot of plant endemism [3]. Within this hotspot, the vine is associated with evergreen forests distributed from 32° S to 40° S, and from 100 to 1000 m above sea level [2, 4]. There is also a record of this species on Juan Fernández Island [5], although this record needs to be confirmed.

This endemic vine is recognizable by its flexible stem and dark green coriaceous leaves which can be bi- or tri-ternate [2, 6, 7]. The vine is dioecious [8]; male plants produce dark violet flowers in pendant clusters (Fig 1A), while female plants produce solitary flowers of the same colour (Fig 1B) [7]. The fruits of the plant are ovoid or oblong, measuring approximately 5.0 cm in their longest axis (Fig 1C) [9, 10]. They have a greenish or yellowish epicarp occasionally covered by a variable number of irregular violet spots [9, 10]. The fruits also have a whitish pulp containing several black seeds [8]. The pulp is edible, fragrant, and sweet, making it highly valued by the locals [7]. Although the fruits are highly sought after, there are no production systems for this plant, and its fruits are still collected via wild harvesting [4].

**Fig 1.**
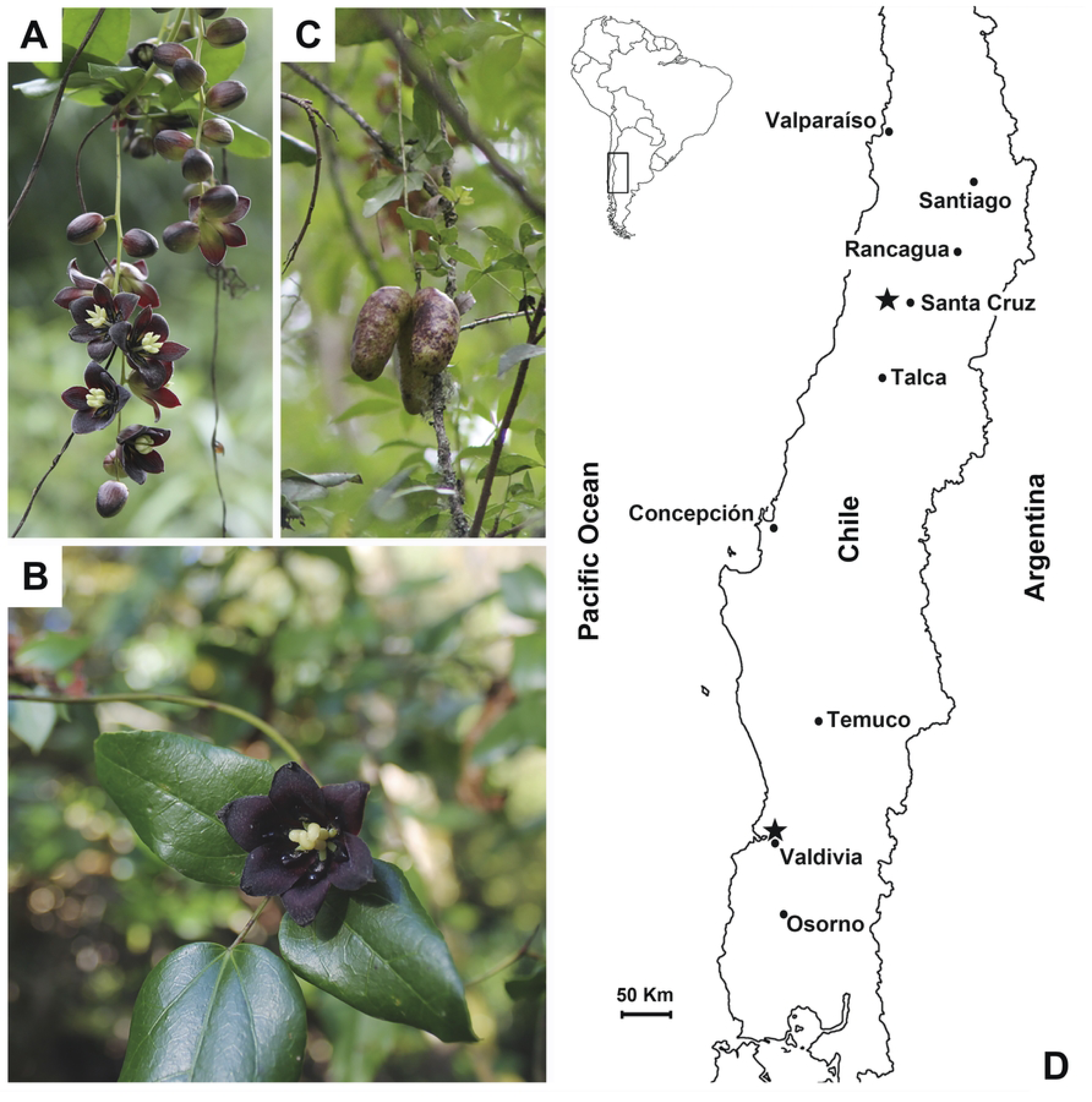
Flowers, fruits, and distribution of *Lardizabala biternata*. A) Male flowers, B) female flower, C) fruits, and D) sampling location of *Lardizabala biternata* fruits. Partial map of central Chile that shows the location of fruit sampling in Valdivia during the 2013, 2016, 2018 and 2019 seasons, and Santa Cruz city during the 2016 season (black stars). In the corner, a map of South America is depicted, and the inside square provides an amplification of the study area.

The scientific knowledge about *L. biternata* is limited, despite botanists having been acquainted with this species for over 300 years [7, 11, 12]. It wasn’t until the 1960s that a method to isolate oleanolic acid from its leaves was developed [13]. Wood anatomy research on this vine took place in the 1980s [14], and its position within the Lardizabalaceae family was only genetically confirmed in the 1990s [15]. Currently, a lack of knowledge persists regarding the ecological role and dispersal mechanism of *L. biternata*, although there are hints that foxes (i.e., *Lycalopex griseus* and *L. culpaeus*) occasionally consume and disperse the fruits and seeds of this vine [16]. For the moment, the information on *L. biternata* is limited and indirect, and these publications do not address plant physiology or the relationship between morphological traits, which are key tools for evaluating and classifying germplasm, as well as for genetic breeding.

In contrast to other vine species belonging to the Lardizabalaceae family [17, 18], there is no morphometric information on the fruits of *L. biternata*. We completely lack knowledge on how its fruits vary in terms of weight, length, seed number, among other morphological traits. This is because a comprehensive morphometric and morphological analysis of the fruits of this endemic and monotypic vine species has never been conducted. The absence of this information has limited informed decision-making regarding variety selection, agronomic practices, and management strategies to optimize the yield and quality of the harvest of this plant [19-26].

In this study we accomplished two goals. First, we investigated qualitatively and quantitatively the morphological traits of *L. biternata* fruits based on the criteria applied in other members of Lardizabalaceae [17, 18, 27]. Second, we explored the relationship between morphological traits of *L. biternata* fruits and their edible pulp content as well as seed weight. This information is crucial to direct the development of efficient production systems aimed at optimizing yield and genetic variability among fruiting plants [28-30].

## Materials and methods

### Study sites and climate conditions

To investigate the phenotypic traits of *L. biternata* fruits we collected fruits of this plant at two contrasting sites in Chile. The first site, Santa Cruz is located in central Chile (34°70’74.9’’S - 71°46’57.7’’W), while the second, Valdivia is located in south-central Chile (39°80’42.8’’S - 73°24’95.7’’W), approximately 500 km south of Santa Cruz (Fig 1D). These sites represent the northern and southern limits of the geographic distribution of *L. biternata* (Fig 1D). Santa Cruz experiences a warm Mediterranean climate, with summer temperatures averaging between 27 and 31 °C [31]. The annual precipitation in this location is around 700 mm, with the majority of rainfall occurring during the winter months (June to August). Valdivia, on the other hand, has an oceanic climate with Mediterranean influences. During the summer, the average temperature is about 17 °C, while in winter, the temperature descends to 8.5 °C. Average annual precipitation is 1,750 mm, distributed throughout the year but concentrated mainly during the winter [32].

### Fruit sampling and climatic data collection

At each study site, we randomly searched for and harvested ripe fruit of *L. biternata*. In Santa Cruz, we collected fruits during the 2016 summer season (StCr, n = 53 fruits), while in Valdivia, we collected fruits during the summer season of 2013 (Vald13, n = 40 fruits), 2016 (Vald16, n = 73), 2018 (Vald18, n = 79) and 2019 (Vald19, n = 37). Harvested fruits were immediately stored in plastic bags and placed inside a cooler with gel ice packs to prevent fruit dehydration and loss of fresh weight. Fruits were transported to the laboratory for investigate of their phenotypic traits.

We obtained annual records of air temperature and rainfall from weather stations located near the study sites because these climatic factors play a crucial role in the development of Chilean fruits [33]. Climatic data covered the period from March to February of next year, which would correspond to the hypothetical biological cycle of *L. biternata*. In Santa Cruz we obtained annual climatic data from the i) El Membrillo (34°80’S – 71°62’W) and ii) El Tambo, San Vicente (34°47’S – 70°99’W) weather stations (from March 2015 to February 2016). In Valdivia, we obtained data from the weather station of the i) Universidad Austral (39°81’S – 73°25’W; from March 2012 to February 2013), and from the weather station of the ii) Estación Experimental Agricultura Austral (39°79’S – 73°23’W; from March 2015 to February 2016, from 2017 to 2018 and from 2018 to 2019).

### Morphological traits of *L. biternata* fruits

We conducted qualitatively and quantitatively analysis on the collected fruits to investigate the morphological traits of *L. biternata* fruits. During the qualitative analysis we recorded the shape and colour of peel, pulp, and seeds. Then, fourteen morphological and morphometric traits were evaluated, including individual fresh weight (IFW), fruit length (FL), fruit width (FW), fruit height (FH), fruit diameter (FD), fruit volume (FV), edible pulp weight (EPW), seed weight (SdW), peel weight (PeW), total seed number per fruit (TSdn°), viable seed number per fruit (VSdn°), non-viable seed number per fruit (FSdn°), individual seed weight (ISdW) and edible pulp plus peel weight (EPW+PeW).

The weight (IFW) and size (FL, FW, FH, FD and FV) of fruits were recorded in all seasons and for all fruits. The IFW of each fruit was measured using an electronic balance (Mettler Toledo, XP205DR, Greifensee, Switzerland). Traits such as FL, FW, and FH were measured on the same fruits using a digital calliper (6-inch 150 mm Digital Calliper, China). The fruit length was measured as the maximum distance between the peduncle and the apex (Fig 2). Both the width and height of fruits were measured using the fruit the longitudinal suture of the fruit as a reference (Fig 2). The width of the fruits was measured perpendicular to the suture, while the height was measured as the distance between the sutures on both sides of the fruit. The diameter of each fruit (FD) was estimated as the average between the width and height of each fruit [FD = (FW × 0.5) + (FH × 0.5)]; while the FV was calculated as that of a cylinder [FV = FL × 3.1416 × (FD × 0.5)^2^].

**Fig 2.**
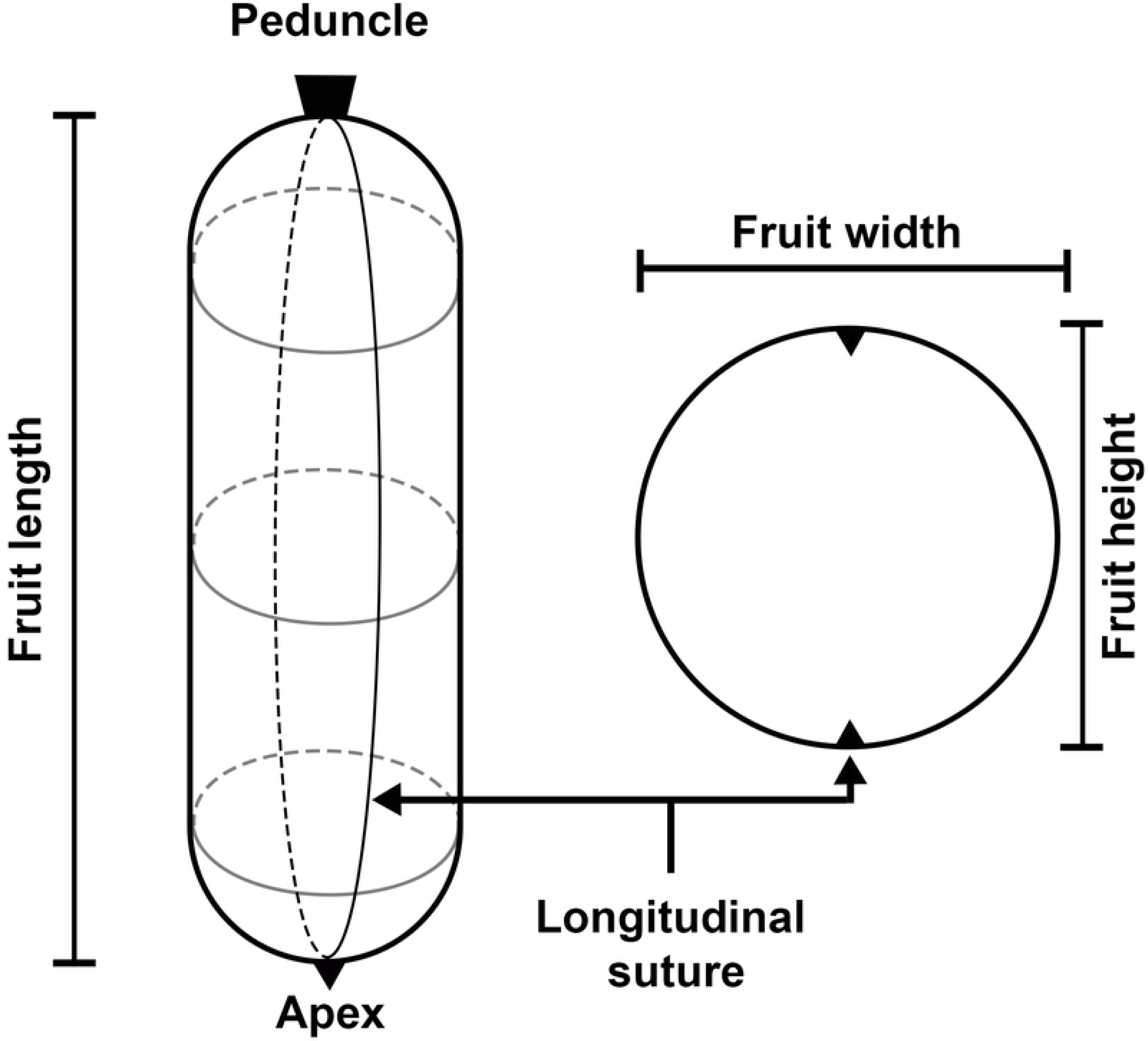
Morphology of *Lardizabala biternata* fruits. Graphic model of *L. biternata* fruits, showing the length, width, and height.

The EPW, SdW, PeW, TSdn°, VSdn°, FSdn°, ISdW, and EPW+PeW were recorded in three subgroups of fruits that were previously measured. Fruits were randomly selected from the Vald16 (20 fruits), Vald18 (10 fruits) and StCr16 (15 fruits) seasons. To estimate PeW, we removed the peel from the fruit using a scalpel. Then, we gently scraped the inner side of the peel with a spoon to remove any remaining of edible pulp, washed it with tap water and gently wiped it dry with tissue paper. We then weighed the peel on an electronic balance.

To estimate seed parameters, we placed the peeled fruit in a metal strainer, washed it under running water, and pressed it with a sponge to remove the pulp and retain the seeds. During washing, we separated and counted futile and viable seeds to estimate FSdn° and VSdn°, respectively. We considered as futile seeds those seeds that floated in the supernatant and as viable those that rested at the bottom of the strainer (S1 Fig). We then summed the values of FSdn° and VSdn° to estimate TSdn°. To estimate SdW, we deposited all seeds on tissue paper, allowed them to air dry for nearly one hour and then weighed them on an electronic balance. Finally, we estimated ISdW as the ratio of SdW to VSdn° (ISdW = SdW / VSdn°).

To estimate the edible pulp weight, it was calculated as the difference between IFW and the sum of SdW and PeW [EPW = IFW – (SdW + PeW)]. To determinate the combined weight of edible peel and pulp of each fruit, we calculated the sum of EPW and PeW [(EPW+PeW) = EPW + PeW], because the peel of *L. biternata* fruits does not register toxicity or contraindications. The percentage of seeds, peel, edible pulp, and peel plus edible pulp were estimated relative to IFW.

### Statistical analysis

We considered each year and location (sampling) as a population. Statistical measures of central tendency and dispersion were estimated for all fruit traits, including the average (mean), maximum value (max), minimum value (min), standard deviation (SD), standard error (SE) and coefficient of variation (CV). One-way analysis of variance (ANOVA) was used to determine de statistic difference between dimensions of *L. biternata* fruits in each population (FL vs FW vs FH), and the differences between fruit components (EPW vs SdW vs PeW).

We conducted an unbalanced analysis of variance [34], because the data number were dissimilar, on each of these statistical measures to investigate whether fruits varied morphometrically between harvesting periods (seasons). When significant effects were detected, a Tukey test (5 %) was applied to evaluate differences among averages. Finally, linear regression analyses were used to evaluate the degree of association between variables. Principal component analysis (PCA) was performed on the traits data from Vald16, Vald18 and StCr16, a biplot was constructed based on the first two principal components (i.e., PC1 and PC2) of the PCA All statistical analyses were conducted using INFOSTAT software in, 2020 [35].

## Results

### Temperatures and precipitation

In Santa Cruz, located to the north of the geographic distribution of *L. biternata*, rainfall accumulated 586 mm between March 2015 and February 2016 (Table 1). Rainfall was not evenly distributed throughout the year, but concentrated between winter and early spring (S2 Fig). The air temperature exhibited annual minimum and maximum averages of 8.8 and 23.1 °C, respectively (Table 1). Occasionally, the air temperature reached 30 °C during the summer (Table 1 and S2 Fig).

**Table 1.**
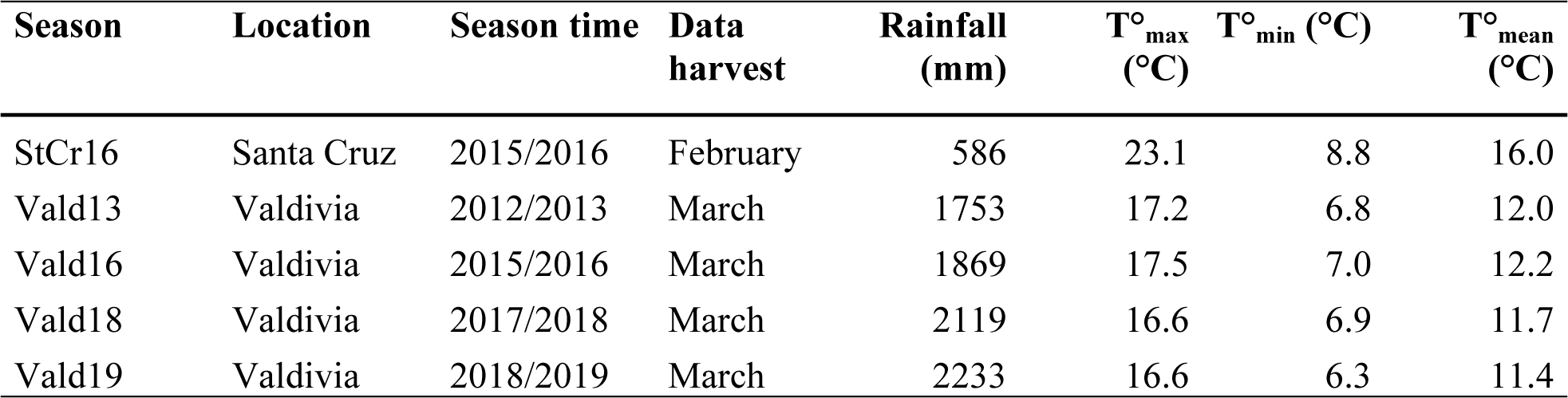
Rainfall and temperatures in Santa Cruz and Valdivia during *L. biternata* fruits growth. **Footnote:** Rainfall accumulates, and average of maximum, minimum and daily temperatures in Santa Cruz and Valdivia Santa Cruz during the growing season (March to February). Data were recorded in Santa Cruz during 2015-16 season (StCr16), and in Valdivia during the 2012-13 (Vald13), 2015-16 (Vald16), 2017-18 (Vald18) and 2018-19 (Vald19) seasons. The data show the rainfall (mm), as well as maximum (T°_max_) minimum (T°_min_) and daily (T°_mean_) average temperatures.

In Valdivia, located to the south of the geographic distribution of *L. biternata*, rainfall was more abundant than in Santa Cruz. Rainfall in Valdivia averaged 1,994 mm per year (Table 1) across four seasons studied (March and February 2012-2013, 2015-2016, 2017-2018 and 2018-2019). Valdivia’s air temperature was lower than in Santa Cruz, averaging over the investigated period a minimum and maximum of 6.8 and 17.0 °C, respectively (Table 1 and S2 Fig).

### Fruit characteristic

Our morphological (qualitative) analysis revealed that most *L. biternata* fruits are roughly ellipsoidal or cylindrical with rounded ends (Fig 3A, 3B and 3C). Usually, one end is slightly wider than the other (Fig 3C). The sides of the fruits are not straight; one is slightly convex and the other slightly concave. Some fruits are cylindrical, although this morphotype is rather unusual (Fig 3A, 3B and 3C). Immature fruits have a smooth and shiny peel, while mature fruits develop large bumps and small granules distributed in patches under the peel that give them a slightly rough texture to the touch (Fig 3C). The peel of the fruits is thin, olive green (immature) and yellow (mature) and is partially or completely covered with light or very dark purple spots of varying size and irregular edges (Fig 3A, 3B and 3C). A longitudinal section of the fruit showed that, inside, it has a shiny white or pale gray pulp surrounding seeds arranged along the entire length of the fruit (Fig 3D). A cross section further revealed that the seeds are arranged radially, forming a circular pattern in the centre of the fruit (Fig 3E). The seeds are large, shiny black and more or less rectangular in shape (Fig 3F). The seeds took on a dark brown colour when extracted and air-dried (Fig 3F).

**Fig 3.**
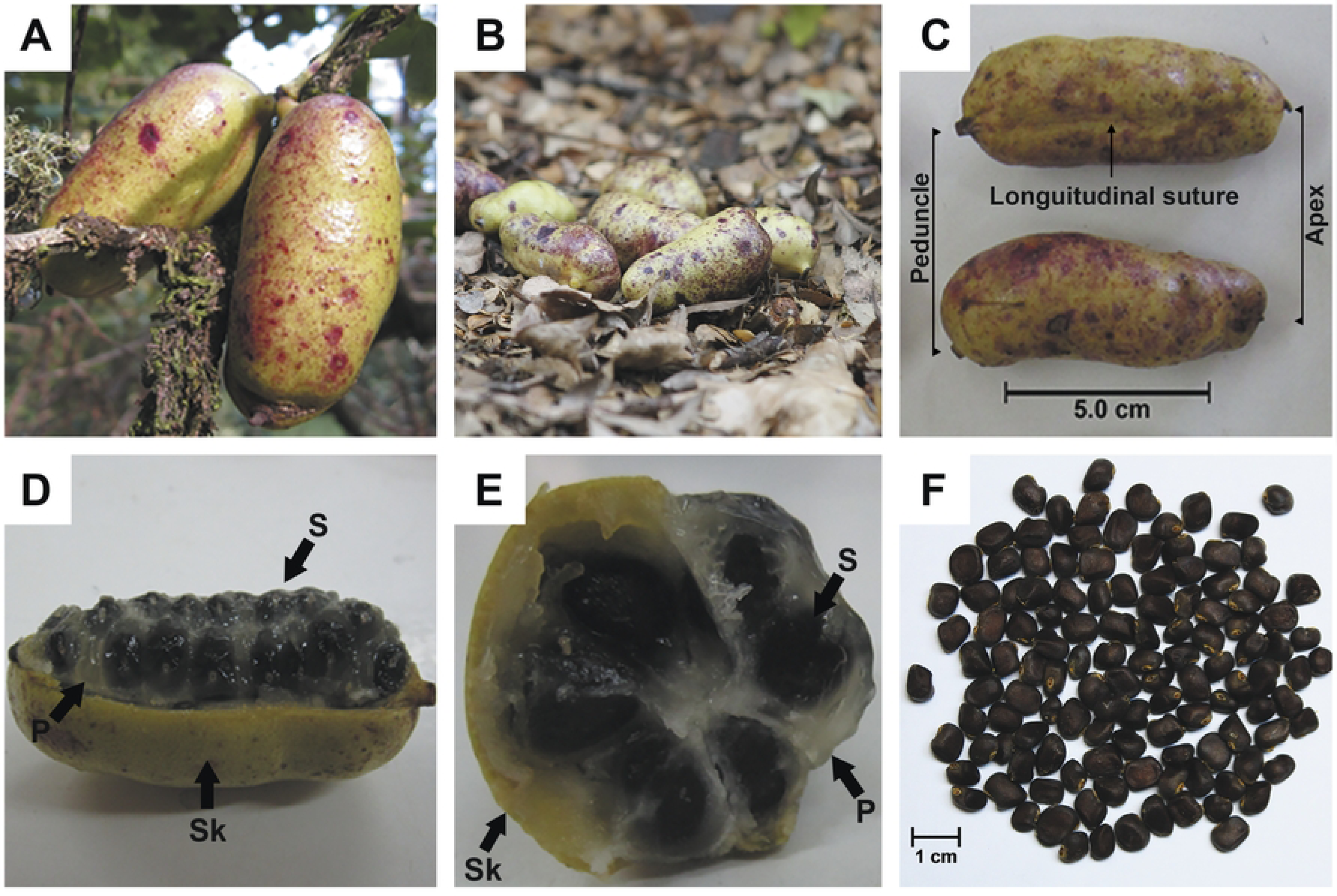
Fruits and seeds of *Lardizabala biternata*. A) Fruits on the plant, B) fruits on the soil, C) harvested fruits, D) fruit dissected in frontal E) and transversal planes, and F) seeds of *L. biternata*. Letters indicate the edible pulp (EP), seeds (Sd) and peel (Pe) of *L. biternata* fruits. Bars represent scales of 5.0 and 1.0 cm.

### Weight and size of *Lardizabala biternata* fruits

Fresh fruits weighed (IFW) between 3.0 and 44.6 g (Table 2), averaging 20.8 ± 0.5 g (mean ± s.e.) (CV = 41.5 %), and exhibiting a normal distribution of data (Figure not shown). Fruits harvested in Santa Cruz reached up to 36.4 g, with an average weight similar to that of fruit harvested during Vald13 and Vald16 (Table 2). Fruits from Vald18 and Vald19 recorded highest weight, being up to 56 % heavier than fruits from Vald13 (P < 0.001).

**Table 2.** Individual fresh weight (IFW) of *Lardizabala biternata* fruits. **Footnote:** Fruits were harvested in location in both Santa Cruz and Valdivia. Data were recorded for fruits from Santa Cruz during 2016 season (StCr16), and from Valdivia during the 2013 (Vald13), 2016 (Vald16), 2018 (Vald18) and 2019 (Vald19) seasons. The data present the average individual fresh weight (mean), minimum (min), maximum (max), standard deviation (SD) and coefficient of variation (CV %). Different letters indicate differences according to Tukey’s test (P < 0.05).

| Season | Individual fresh weight of fruits (g) |  |  |  |  |  |
| --- | --- | --- | --- | --- | --- | --- |
|  | Fruits (n°) | Mean (g) | Min (g) | Max (g) | SD (g) | CV (%) |
| StCr16 | 53 | 18.0<sup>b</sup> | 3.0 | 36.4 | 8.6 | 47.6 |
| Vald13 | 40 | 16.6<sup>b</sup> | 3.8 | 27.9 | 6.0 | 36.1 |
| Vald16 | 73 | 17.9<sup>b</sup> | 3.9 | 35.0 | 7.9 | 44.1 |
| Vald18 | 79 | 25.9<sup>a</sup> | 5.4 | 44.6 | 8.5 | 32.7 |
| Vald19 | 37 | 24.3<sup>a</sup> | 13.6 | 38.2 | 6.4 | 26.2 |
| Total |  | 20.8 | 3.0 | 44.6 | 8.6 | 41.5 |

Fruit morphology showed contrasting differences between dimensions (FL vs FW vs FH), as well as in the dispersion of data (Fig 4). For instance, the FL recorded higher values than both width or height in the same fruits (P < 0.001), while FW and FH showed similar values between them (Fig 4). The FL displayed a wide range of variation, measured between 20.1 and 83.4 mm in length, with an average of 54.0 ± 0.8 mm (CV = 24.1 %) (Fig 4A). The FW and FH averaged 24.4 ± 0.2 mm (CV = 12.4 %) and 24.4 ± 0.2 mm (CV = 12.6 %), respectively (Fig 4B and 4C). Fruits were influenced by the season (P < 0.001), Vald18 fruits were 17.4 % longer than StCr16 fruits (Fig 4A), and 13.4 % wider than Vald13 fruits (Fig 4B). The FD and FV averaged 23.7 ± 0.2 mm (CV = 12.2 %) and 24,949 ± 579 mm^3^ (CV = 39.0 %), respectively. Both characters were affected by the season (P < 0.001), e.g. fruits harvested in Vald19 were up to 33.5 % more voluminous (Figure not shown).

**Fig 4.**
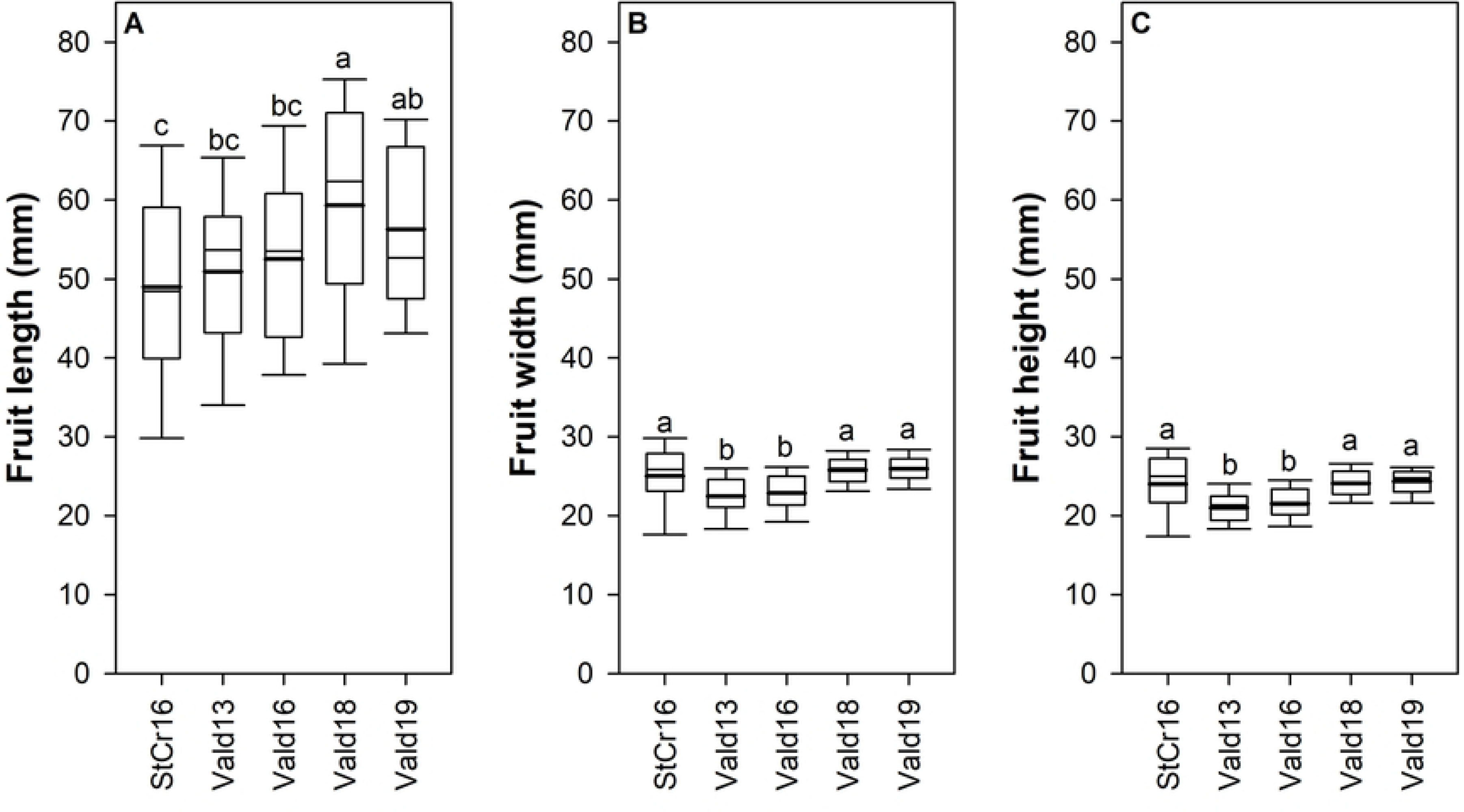
Size of *Lardizabala biternata* fruits. A) Length, B) width and C) height of *L. biternata* fruits from Santa Cruz during the 2016 season (StCr16), and Valdivia during the 2013 (Vald13), 2016 (Vald16), 2018 (Vald18), and 2019 (Vald19) seasons. Box-and-whisker plots depict the morphometric parameters; the lower limit of the box indicates the 25th percentile, the black line represents the median (50th percentile), and the upper limit of the box indicates the 75th percentile. The error bars on either side of the box indicate the 10th and 90th percentiles. The black line within the box marks the mean, and different letters indicate significant differences between populations after unbalanced ANOVA and Tukey’s test (P < 0.05).

The IFW showed a positive association with the length (Fig 5A), width (Y = 2.10X – 30.62; R^2^: 0.54 and P < 0.001), and height (Y = 2.13X – 28.11; R^2^: 0.51 and P < 0.001) of the same fruits. Additionally, the IFW registered positive association with parameters calculated as the diameter (Y = 2.23X – 32.21; R^2^: 0.56 and P < 0.001) and the volume of the fruits (Fig 5B). The highest coefficients of determination were observed between IFW vs FL and IFW vs FV (Fig 5).

**Fig 5.**
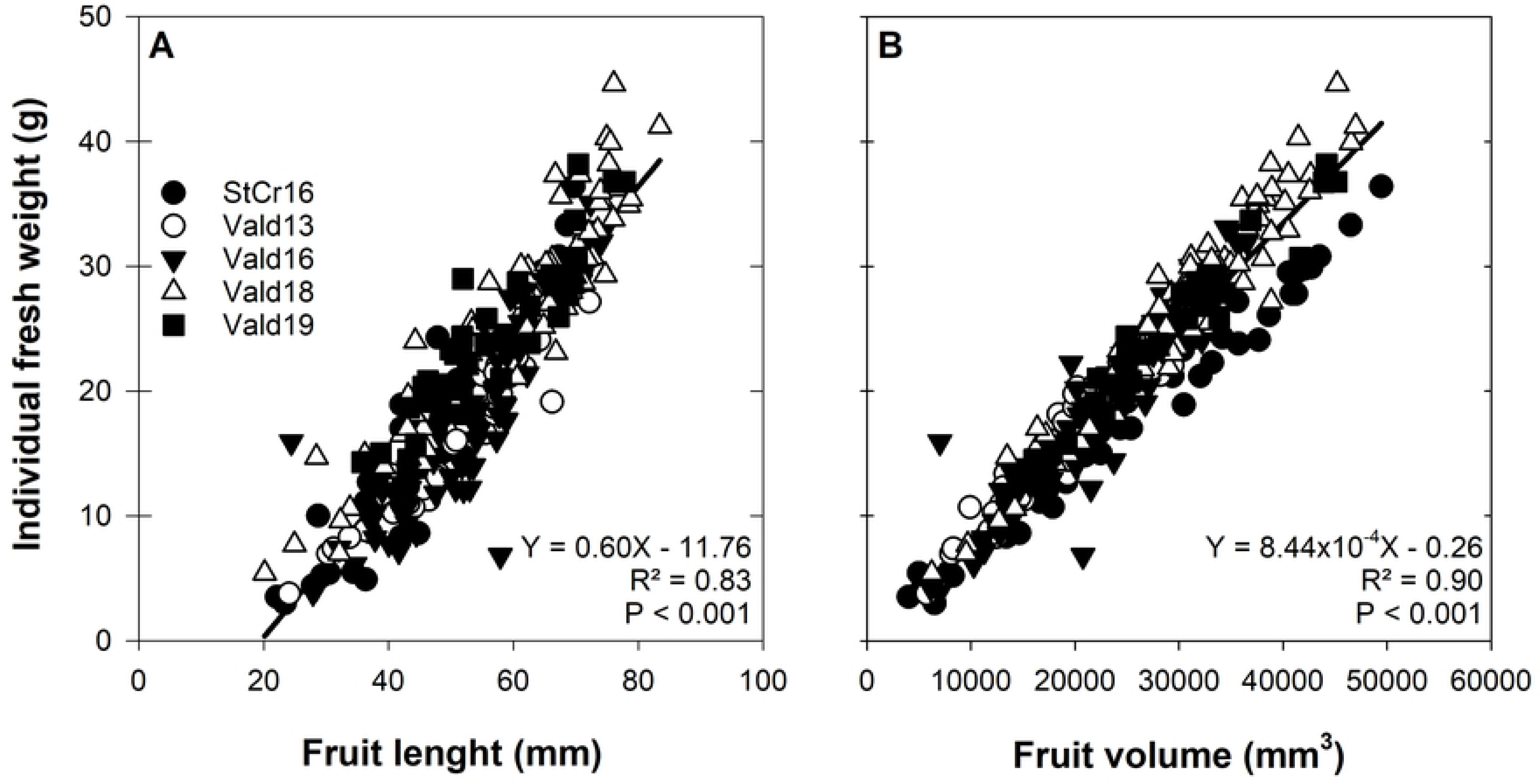
Association of morphological traits of *Lardizabala biternata* fruits. A) Association of the individual fruits weight (IFW) with the fruit length (FL), and B) fruit volume (FV) of *L. biternata* fruits from the Santa Cruz 2016 (StCr16) season, and from Valdivia during the 2016 (Vald16), 2018 (Vald18) and 2019 (Vald19) seasons. Solid lines represent the regression analysis.

### Weight of edible pulp, seeds, and peel of *Lardizabala biternata* fruits

Subgroups of fruits weighed 25.7 g (11.9 – 36.4 g), 20.0 g (12.1 – 30.0), and 29.6 g (18.1 – 39.9 g) in StCr16, Vald16, and Vald18, respectively. Previous fruits registered statistics differences between seasons (P < 0.01). Fresh weight of the pulp (EPW), seed (SdW) and peel (PeW) exhibited considerable variation (Fig 6), registering statistics differences between them (P < 0.071). For instance, EPW averaged 10.5 ± 0.5 g (CV = 30.1 %), and it was main contributor of the IFW (34.9 to 53.7 %) (Fig 6). In the same way, seeds inside *L. biternata* fruits weighted between 1.6 and 13.6 g to IFW, contributing from 13.4 to 40.3 % to the IFW (Fig 6). The peel covering the *L. biternata* fruits weighted from 1.7 to 10.2 g, contributing between 12.7 and 43.7 % of the IFW (Fig 6). Regarding sampling, fruits harvested in Vald16 registered higher percentage of EPW (P < 0.001), at the same time, a less percentage of PeW (P < 0.001). The seed percentage did not show a difference between locations (P = ns), contributing to 30.2 % of the IFW (CV = 18. 4 %). The sum of EPW and PeW (EPW+PeW) averaged 16.7 ± 0.8 g (min = 8.2 g and max = 27.0 g), while the CV of 31.5 % (data not shown). EPW content was not affected by seasons (P = ns).

**Fig 6.**
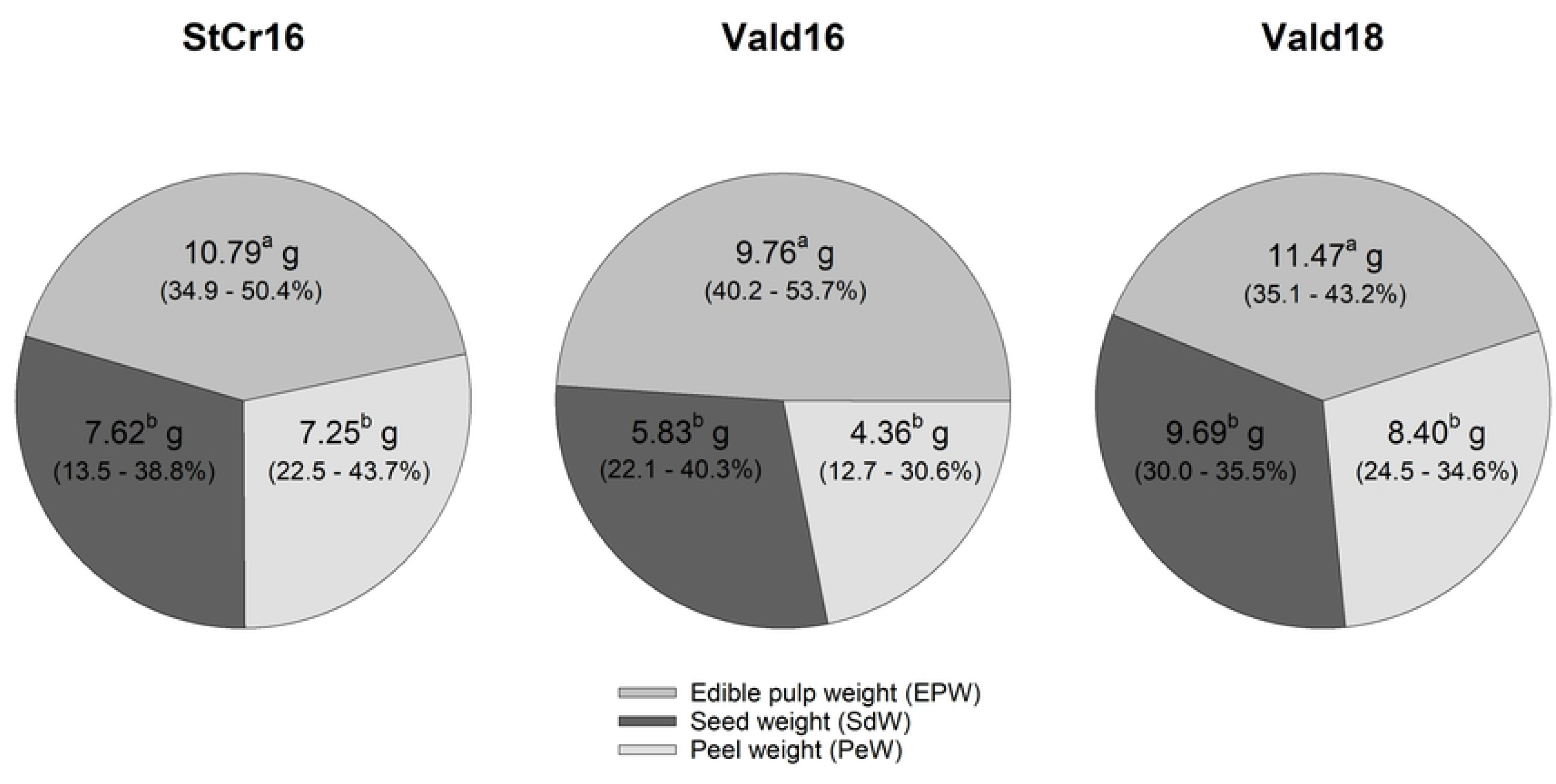
Weight of edible pulp, seeds, and peel of *Lardizabala biternata* fruits. Average weight of edible pulp (EPW), seeds (SdW), and peel (PeW) of *L. biternata* fruits from the Santa Cruz 2016 season (StCr16), and from Valdivia 2016 (Vald16), and 2018 (Vald18) seasons. Contributions of EPW, SdW, and PeW to individual fresh weight in percentage. Different letters indicate significant differences between fruit components after ANOVA and Tukey’s test (P < 0.05).

The individual fresh weight of *L. biternata* fruits showed a positive association with the edible pulp (Fig 7A), seeds (Fig 7B), and peel (Fig 7C) of the same fruits. Additionally, the length, width and height of the fruits exhibited a positive association with the weight of internal structures of the same fruits (data not shown). The edible pulp weight registered a positive association with the seed weight (Fig 7D) and peel weight (Fig 7E) of the same fruits, as well as with fruit length (7F) and other morphological traits (Data not shown).

**Fig 7.**
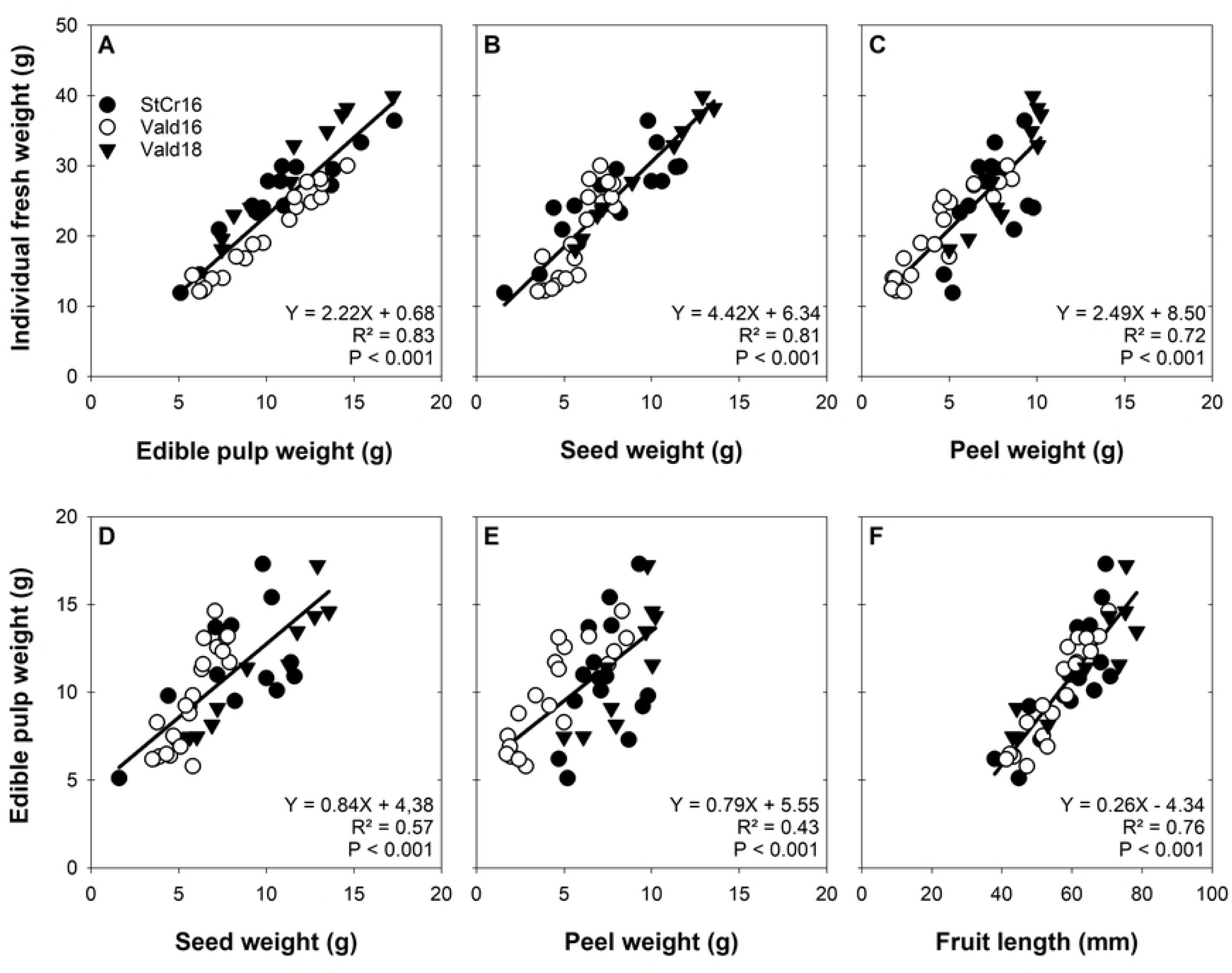
Association of internal structures of *Lardizabala biternata* fruits. Association of the individual fruits weight (IFW) with the A) edible pulp weight (EPW), B) seed weight (SdW) and C) peel weight (PeW). Simultaneously, the association of the EPW with the D) SdW, E) the PeW and F) fruit length (FL) of *L. biternata* fruits harvested in Santa Cruz during the 2016 season (StCr16), and Valdivia during 2016 (Vald16) and 2018 (Vald18) seasons. Solid lines show the regression analysis.

### Number and weight of seeds of *Lardizabala biternata* fruits

*Lardizabala biternata* fruits showed several seeds along the same fruit, but only 6 seeds were observed in the transverse plane (Fig 3E). On average, fruits contained on 48 ± 2 seeds (TSdn°; 20 to 70 seeds), where 93.1 % were viable (VSdn°) and 6.9 % non-viable (FSdn°) (Table 3). Seed number character did not show statistical difference between seasons (P = ns). The ISdW ranged from 94.0 to 240.2 mg (average =163.7 ± 5.8 mg and CV = 23.5 %), with seeds from Vald18 being 23.3 % heavier than the average seed (P < 0.001).

**Table 3.** Descriptive analysis of seed traits of *Lardizabala biternata* fruits. **Footnote:** Fruits were harvested in Santa Cruz and Valdivia locations. Data were recorded for fruits from the Santa Cruz 2016 season (StCr16), and from the Valdivia 2016 (Vald16), and 2019 (Vald19) seasons. TSdn°: total seed number per fruit; VSdn°: viable seed number per fruit; FSdn°: non-viable seed number per fruit; and ISdW: individual seed weight. Data shows the average (mean), minimum (min), maximum (max), standard deviation of mean (SD) and coefficient of variations (CV %). Different letters indicate Tukey’s test differences (P < 0.05) between seasons.

| Fruit traits | Season | Mean | Min | Max | SD | CV (%) |
| --- | --- | --- | --- | --- | --- | --- |
| Total seeds number per fruit (n°)<br>(TSdn°) | StCr16 | 47,5<sup>a</sup> | 11,2 | 23,6 | 20,0 | 60,0 |
|  | Vald16 | 46,5<sup>a</sup> | 11,3 | 24,4 | 28,0 | 68,0 |
|  | Vald18 | 50,5<sup>a</sup> | 17,2 | 34,1 | 30,0 | 71,0 |
| Viable seeds number per fruit (n°)<br>(VSdn°) | StCr16 | 43,1<sup>a</sup> | 14,6 | 34,0 | 12,0 | 58,0 |
|  | Vald16 | 43,6<sup>a</sup> | 10,6 | 24,3 | 25,0 | 63,0 |
|  | Vald18 | 49,1<sup>a</sup> | 17,7 | 36,0 | 25,0 | 70,0 |
| Non-viable seeds number per fruit (n°)<br>(FSdn°) | StCr16 | 4,4<sup>a</sup> | 7,3 | 166,1 | 0,0 | 27,0 |
|  | Vald16 | 2,9<sup>a</sup> | 2,8 | 99,4 | 0,0 | 11,0 |
|  | Vald18 | 1,4<sup>a</sup> | 2,1 | 151,3 | 0,0 | 7,0 |
| Individual seed weight (mg)<br>(ISdW) | StCr16 | 175,4<sup>b</sup> | 34,4 | 19,6 | 98,3 | 232,6 |
|  | Vald16 | 135,9<sup>c</sup> | 25,6 | 18,8 | 94,0 | 194,6 |
|  | Vald18 | 201,8<sup>a</sup> | 21,2 | 10,5 | 164,7 | 240,2 |

The individual fresh weight of fruits showed a positive association with total seed number (Y = 1.24X + 18.06; R^2^: 0.57 and P < 0.001), as well as with the viable seed number (Y = 1.40X + 11.04; R^2^: 0.52 and P < 0.001) within of the same fruits. However, the association between IFW and ISdW registered a lower coefficient of determination (Y = 2.14X + 112.4; R^2^: 0.18 and P < 0.05). The edible pulp registered a positive association with both seed number (TSdn°: Y = 0.19X + 1.22; R^2^: 0.60 and P < 0.001 and VSdn°: Y = 0.18X + 2.33; R^2^: 0.62 and P < 0.001), but showed an insignificant association with ISdW (Y = 0.016X + 7.82; R^2^: 0.04 and P = ns).

### Correlations between the characters and principal components analysis

Simple correlation coefficient analysis showed significant association between individual fresh weight and other morphological traits (Figs 5, 7 and 8). These associations were able to reach coefficient of correlation greatest to 0.90 (Fig 8), although there were association with few coefficients of correlation such as VSdn° (Fig 8). The association between morphological and morphometric traits registered significant correlations, e.g. FL vs VSdn° or SD vs ISdW (Fig 8), but in some cases, the association were insignificant, such as FL vs ISdW or FH vs TSdn° (Fig 8).

**Fig 8.**
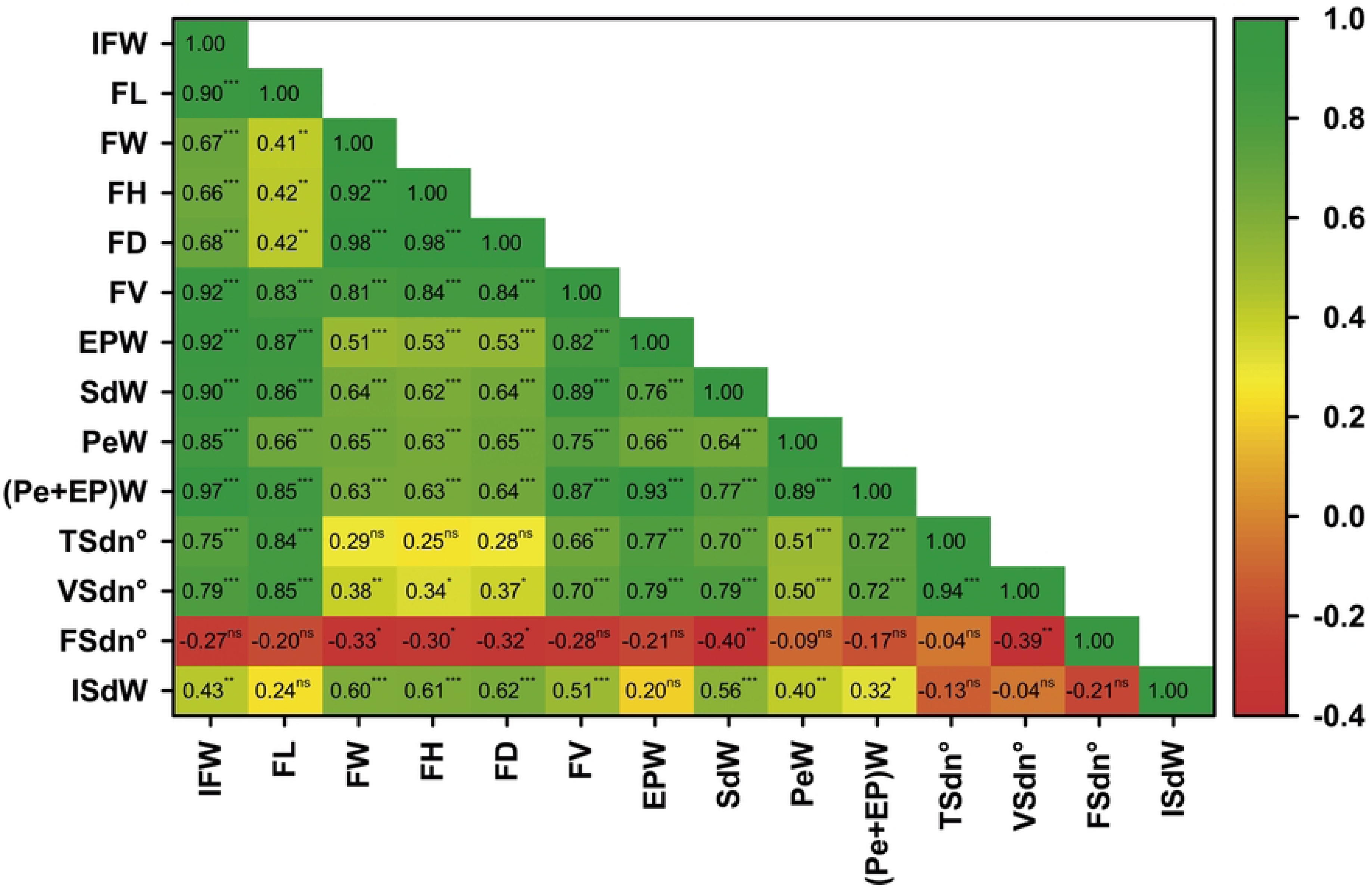
Correlation heat map showing the relationship between morphological traits of *Lardizabala biternata* fruits. The heat map displays the correlation coefficient values between pairs of traits. Data were registered for fruits of *L. biternata* from the Santa Cruz 2016 season (StCr16), as well as from Valdivia 2016 (Vald16), and 2018 (Vald18) seasons. IFW: Individual fresh weight; FL: fruit length; FW: fruit width; FH: fruit height; FD: fruit diameter; FV: fruit volume; EPW: edible pulp weight; SdW: seed weight; PeW: peel weight; EPW+PeW: edible pulp plus peel weight; TSdn°: total seed number per fruit; VSdn°: viable seed number per fruit; FSdn°: non-viable seed number per fruit; and ISdW: individual seed weight. n.s., not significantly different at P < 0.05; *, ** and *** different at P < 0.05, < 0.01 and < 0.001, respectively.

The PCA revealed patterns of variability among phenotypic characteristics (S1 Table). In the analysis, eigenvalues greater than 1.0 were used to decide which principal components (PCs) to include, identifying three principal components that explained 89.2% of the total variation (S3 Fig). Of the first two components, which accounted for 81.7 % of the total variability, PC1 explained 65.0% of the variance, followed by PC2 with 16.7%. The first component consisted of fruit character related to weight (IFW, EPW, SdW and other) and fruit size (FL and FV) (Fig 9), while the PC2 was influenced by characters associated with the transversal growth of the same fruits such as FH, FW, FD and ISdW (Fig 9). PC3 explained the non-viable seed number and accounted for 7.6 % of the variance (S3 Fig).

**Fig 9.**
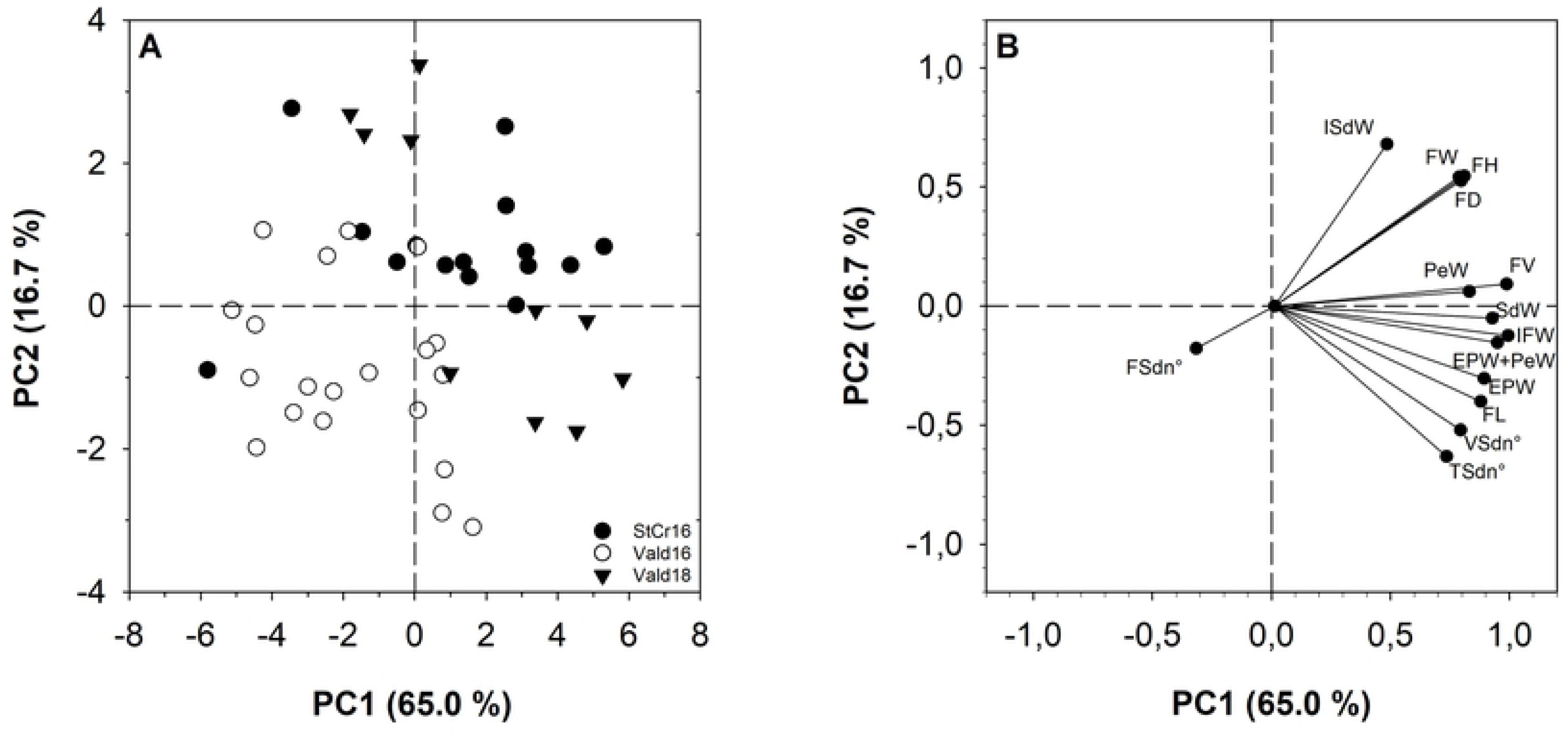
Principal component analysis of physiological traits of *Lardizabala biternata* fruits from Santa Cruz 2016 (StCr16) and Valdivia 2016 (Vald16) and 2018 (Vald18) in the PC1, PC2 plane: score plot (A) and correlation loadings (B). IFW: Individual fresh weight; FL: fruit length; FW: fruit width; FH: fruit height; FD: fruit diameter; FV: fruit volume; EPW: edible pulp weight; SdW: seed weight; PeW: peel weight; EPW+PeW: edible pulp plus peel weight; TSdn°: total seed number per fruit; VSdn°: viable seed number per fruit; FSdn°: non-viable seed number per fruit; and ISdW: individual seed weight.

To identify variations in the selected seasons, as well as phenotypic traits (Fig 9), the bi-plot revealed that the IFW had high positive correlations with SdW, EPW+PeW and SdW, while the EPW showed a better correlation with FL and VSdn° (Fig 9B). Physiological traits that defined transversal growth such as FW, FH and FD, exhibited positive correlations with each other, in addition with ISdW. However, ISdW displayed negative correlations with fruit traits as VSdn° and TSdn° (Fig 9).

## Discussion

Before this investigation, *L. biternata* could be defined as a scientifically unknown plant, lacking domestication and with wasted edible fruits, contrasting sharply with studies observed in seven genera of Lardizabalaceae from East Asia [4]. The quantification of both morphological and morphometric traits of *Lardizabala biternata* revealed in this study is fundamental as the first step for the evaluation and classification of germplasm [27]. But the functionality of morphological traits is much broader; it is an important tool to understand plant physiology, identify the morphological traits that control plant development and growth, analyse the climatological variables that affect the plant cycle, differentiate elite genotypes, or parameterize agricultural technologies [27, 36-39].

### Weight of *L. biternata* fruits

The fruit weight should be one of the primary aims of plant breeding, and the present study revealed that fresh *L. biternata* fruits average 20.8 g in the two selected populations. These fruits exhibit wide variability (41.5 %), with some fruits reaching up to 36.4 g. But, they are lighter that Asian species than have more extensively researched (Table 2), such as *Dicaisnea insignis* (31,4 g) and *Akebia quinata* (72.1 g) [17, 18]. The difference in fruit weights is more significant when comparing the current *L biternata* fruits with improved *A. quinata* fruits under favourable cultivation conditions, which can reach up to 546 g [17, 18]. At present, the maximum fresh weight of *L. biternata* fruits is unknown, as are the favourable conditions to achieve this potential weight. Therefore, fruit breeding aimed at increasing fresh weight, as well as identifying the nutritional conditions necessary to achieve this goal, could be interesting objectives for future research.

In *L. biternata*, each fruit consists of three distinct structures (pulp, seeds, and peel), all which contribute to the IFW (Fig 3). In this instance, the pulp is the primary component of the fruits, contributing over 50 % of fresh weight (Fig 6) and water content (data not shown). While the peel and seeds of *L. biternata* fruits contribute less to the fresh weight (Fig 6), but they have the highest dry matter content among the same fruits (data not shown). The edible pulp content in *L. biternata* is relatively similar to that registered in Asian species, and the seed weight is slightly higher. However, the peel shows contrasting characteristics compared to improved species [17, 18]. For example, the peel of *L. biternata* fruits is thin and light (< 24.5 %), while the peel of *A. quinata* fruits is thicker and heavier, contributing up to 62.9 % of the total dry matter and higher percentages as fresh weight [17, 18].

Undoubtably, in order to propose *L. biternata* fruits or others as food, it is necessary to understand their edible proportion. In this research, the ratio between edible and inedible content can reach over 60 % in *L. biternata* fruits, a proportion higher than that reported in related species such as *A. quinata* (17 to 40 %) and D. *insignis* (48.5 %) [17, 18, 40]. The second quality of *L. biternata* as edible fruit is its peel; to date, there are no medical contraindications or food limitations associated with the consumption of the peel of *L. biternata* fruits in the case of accidental ingestion. If the peel of *L. biternata* fruits is integrated into the edible portion, it would increase the ratio between edible and inedible parts, furthermore, favouring the Chilean fruit compared to the Asian species. Enhancing fruit size and increasing the proportion of edible content are primary objectives of breeding programs. Integrating fruit peel into these programs could streamline improvement strategies. Moreover, studies dedicated to peel characteristics could identify fruit qualities that would guide *L. biternata* fruit potential in the market.

While this study has focused on the potential food use of *L. biternata* fruits, other research could direct their efforts towards the seeds as a source of dry matter, fatty acids, or chemical compounds with pharmacological activity [41-43]. In this research, the total weight, the number, and the individual weight of seeds from *L. biternata* fruits exhibit a wide variability. Furthermore, theses seed traits show associations with traits related to the weight and size of *L. biternata* fruits. Therefore, seeds traits could play a key role in controlling fruit size, as well as influencing seed properties. Theses associations are important because it can pave the way for plant breeding focused on fruit seeds, predict seed content without the need to destroy the biological material, and select fruits with a higher content of edible pulp based on a more permanent trait, among other benefits. However, information about the seeds of *L. biternata* is even more scarce, and it should be further explored in topics such as morphological, biochemical, and nutritional characteristics, as well as conservation and germination aspects.

### Morphological traits of *Lardizabala biternata*

So far, fruit length is the only physiological trait quantified in *L. biternata* fruits [9], but the average fruit length (50.0 mm) could be questioned due to the lack of research defining the morphological traits in the fruits. Our research clarifies the average length of *L. biternata* fruits as well as the complexity the same morphological trait (Fig 4A). For example, i) the average length fruit (52.8 mm) is longer that registered in previous studies (50.0 mm), ii) several fruits measure above the average fruit length (> 45 %), with some even exceeding 70.0 mm (Fig 4A), and iii) some fruits show minimal differences between their length, width, and height (Fig 4). Interestingly, the longest *L. biternata* fruits studied can reach similar lengths to those of Asian species; even so, the average fruit length shows considerable differences, with this endemic fruit being shorter than *A. quinata* (< 104 mm) or *D.insignis* (< 111 mm) fruits [17, 18]. Nevertheless, our study demonstrates that fruit length traits could be fundamental in understanding the physiology of *L. biternata* fruits and serve as a key tool in breeding programs. For example, i) the fruit length trait is dynamic over time (data not shown) and exhibits wide variability at harvest, ii) it is morphological trait easy to measure during the growth *in-situ*, without the need to harvest the fruit or destroy the fruits, and iii) this trait is linked to other morphological traits of fruit yield such as EPW, PeW, as well as, weight and number of seeds (Fig 8). Therefore, deepening our knowledge about traits that control the fruit length would be indispensable for achieving a *L. biternata* fruit with desirable agricultural conditions.

Both the width and height of *L. biternata* fruits exhibit lower values and narrower variability compared to the fruit length of the same fruits (Fig 4). Furthermore, there are few differences between the width and height of fruits, and combining both traits as diameter or estimating the fruit volume could facilitate the comparative study of fruits [44-48]. In *L. biternata*, there are not records regarding width and height of the fruits, and our study demonstrate that both traits are important tools for analysing variability between season and locations (Figs 4B and 4C). Additionally, they serve as the mathematical basis for estimating the diameter (FD) and volume (FV) of the same fruits. These traits are often used to understand physiological control in fruits [44-48], as well as the environmental impact on the same fruits [49-52]. For now, width and height should not be discarded as morphological traits for analysing *L. biternata* fruits, and further studies could reinforce their genetic diversity, environmental variability, and productive importance.

Volumetrically, *L. biternata* fruits are considered as oblong or suboblong (Fig 3A, 3B and 3C), and our research reinforce the contrasting differences between FL versus both FW and FH (Fig 4), but some fruits are shorter, and the differences between dimensions are minimal (Fig 4). The main variability in fruit size is due to the length of the same fruits, which increases the volume and widens the gap with respect to width and height (Fig 4). As the fruit length shows a close association with the weight and number of seed, an increase in the number of viable seeds along the fruits results in an increase in the length of the same fruit. Unlike of fruit length, the width, height, and diameter of fruits are more consistent traits and exhibit less variability between them or between fruits (Fig 4). In response, our research displays the seed organization along the transverse axis of *L. biternata* fruits, showing six seeds growing radially from the centre of the fruit to the peel (Fig 3E). Putatively, the individual weight and size of the seeds should be the main traits that control the transverse growth of *L. biternata* fruits, thereby limiting the variability of the width, height, and diameter of the same fruits. Therefore, if the plant breeding of *L. biternata* fruits is focused on seeds traits, this strategy could take two approaches, i) increasing the seed number via fruit length, or ii) increase the seed weight via fruit diameter. Nevertheless, it is necessary to continue delving into the traits that control the size of L. biternata fruits, as well as incorporating new traits that could have a profound impact on growth, such as growth dynamics, water volume, number of ovules, seed size, among others, to develop agricultural strategies for genetic breeding and maximize fruit size.

### Impact of climatic conditions

The distribution area of *L. biternata* is wide latitudinally, resulting in contrasting climatic permeameters [32, 53, 54]. Our research reinforces the differences in both rainfall and temperatures in the two identified locations, but climatic distinctions being less pronounced between the seasons studied in Valdivia (Table 1). Commonly, the Santa Cruz location experiences low rainfall and high temperatures, but the season studied saw spring rainfall, which are infrequent (S2 Fig). These climatic conditions could affect the genotype per ambient interaction, increasing the variation range of IFW and size of *L. biternata* fruits, thereby potentially overshadowing the negative effect of low rainfall and hight temperatures. If focused the study on Valdivia location, morphologic traits as the IFW and fruit size exhibit sensitivity to climate, as evidenced by the fact that fruits harvested during the Vald18 and Vald19 seasons are both heavier and larger (Table 1 and Fig 4). Both seasons records higher rainfall and lower temperatures compared to the Vald13 and Val16 seasons (Table 1).

Previous studies have shown that low rainfall and high temperatures, similar to those in central Chile, could reduce water sources and increase evapotranspiration [55], which would have a negative effect on the size and weight of *L. biternata* fruits from StCr16, Vald13, and Vald16. However, the small number of populations analysed could obscure the genetic diversity and phenotypic response of *L. biternata*, such as drought resistance, pollination vectors, duration of flowering, and other factors [56]; considering multiples ecoregions in Chile based on weather conditions and elevation above sea level, that have not been studied either

### Correlations between the phenotypic characters

Defining the relationship between traits is indispensable for *L. biternata* breeding programs. For example, it guides appropriate selection, evaluate the influence between them or indirectly estimates the target traits via secondary trait with high association [36, 57]. Our research shows that the fresh weight and pulp content of *L. biternata* fruits exhibit a positive association between them. At the same time both traits registered positive association with FL, FW, FH, FD, FV, EPW, SdW, PeW and VSdn° (Fig 8), indicating that putative target traits would evolution in the same direction as another trait [39]. Furthermore, as the study resolute a positive correlation between IFW and EPW, it would allow to select genotypes with larger fruit size and higher edible proportion simultaneously [38].

In this research, the IFW increased to a similar extent as SdW, TSdn° and VSdn°, while other studied Lardizabalaceae species have shown a negative relationship between the fresh weight of fruits and seed weight or percentage of edible content [27, 38, 39]. However, ISdW has a weaker corelation with IFW and is not significantly related to EPW (Fig 8). Possibly, the longitudinal number of seed could control both the weight and pulp content of the same fruits, while the radial number, size and organization of these seeds could define the transverse size of *L. biternata* fruits. The balance between numerical and gravimetric traits registered in seeds would be necessary to achieve bigger fruits with fewer seeds. This criterion is highly valued because consumers are reluctant to eat fruits with a high number of seeds, as they require spitting them out [27, 38, 39].

The present principal component analysis explains the traits in the form of factors, reducing the number of effective discriminating parameters and suggesting genetic linkage between controlling characters through association among them [58]. In this case, individuals in the lower right corner of the plot were characterized by larger fruit size and higher pulp content. However, selected candidate plants should be tested under different climate conditions to maximize the pulp content.

## Conclusions

This research records fourteen morphological traits of *L. biternata* fruits, similar to traits studied in other Lardizabalaceae species. These morphometric, gravimetric, and numeric traits exhibited different degrees of phenotypic variation, and their average values can characterize *L. biternata* fruits. For example, the fruit weighed 20.8 g and measured 54.8 mm in length and had a diameter of 23.7 mm. Furthermore, the content of edible pulp was 10.5 g, contributing to 44.4% of the total weight of the fruit. Statistical analysis showed variances in fruit traits, as well as differences between traits (FL vs FW vs FH, and EPW vs SdW vs PeW). The correlation analysis indicated the presence of positive or negative correlations among the 14 morphological traits. In this sense, fruit weight showed positive correlations with most fruit traits, such as edible pulp weight, fruit length, as well as the number and weight of seeds, while the diameter of fruits (width and height) showed a higher association with individual seed weight. Our results provided essential information about *Lardizabala biternata* fruits, including both the variability of morphological traits and the association among traits. Furthermore, this information provides genetic parameters and some novel insights into the selection of elite genotypes, genetic management strategies, physiological tools to use, and conservation of biological material. It especially helps to dispel ignorance about the edible fruits of *Lardizabala biternata*.

## Acknowledgments

The first author thanks to his mother Maria Paredes since she was the first reference regarding the fruits of *Lardizabala biternata*; Pablo Ramirez, friend and guide who helped me find the populations of *L. biternata*; Beatriz Shibar, Chief Technician of the Laboratorio de Fitotecnia at the Instituto de Producción y Sanidad Vegetal (Universidad Austral de Chile), where *L. biternata* fruits were analysed; and the staff of Centro de Investigación en Recursos Naturales y Sustentabilidad (CIRENYS) at the Universidad Bernardo ÓHiggins, where this publication was finalized. This study was partially funded by ANID FONDECYT regular 1220605 project awarded to LDF and Dirección de Investigación, Vicerrectoría de Investigación y Postgrado, Universidad de Las Américas.

## Supporting information

**S1 Fig. Seeds of *Lardizabala biternata* fruits.** A) Viable and B) non-viable seeds.

(TIF)

**S2 Fig. Rainfalls and air temperatures.** Accumulated rainfalls (grey bars), and minimum (close triangle plus black dotted line), maximum (close circle plus black line), and mean temperature (open circle plus dotted line) from March to February in Valdivia city during 2012-13 (A, Vald13), 2015-16 (B, Vald16), 2017-18 (C, Vald18), 2018-19 (D, Vald19) seasons and close Santa Cruz city during 2015-16 (E, StCr16) season.

(TIF)

**S3 Fig. Variance percentage accounted by three principal components in PCA for the physiological characters in *Lardizabala biternata* fruits**.

(TIF)

**S1 Table. Eigenvectors of principal component axes from PCA for the physiological characters in *Lardizabala biternata* fruits.**

(DOCX)

